# Transport of Ketoprofen in Mammalian Blood Plasma

**DOI:** 10.1101/2021.04.03.438117

**Authors:** Mateusz P. Czub, Ivan G. Shabalin, Wladek Minor

**Affiliations:** Department of Molecular Physiology and Biological Physics, University of Virginia, 1340 Jefferson Park Avenue, Charlottesville, VA 22908, USA; Center for Structural Genomics of Infectious Diseases (CSGID), University of Virginia, 1340 Jefferson Park Avenue, Charlottesville, VA 22908, USA

**Author notes:** Laboratory of Biomolecular Research, Paul Scherrer Institute, Forschungsstrasse 111, 5232 Villigen, Switzerland (Mateusz P. Czub). Lead Contact: Wladek Minor.

**Keywords:** ketoprofen, non-steroidal anti-inflammatory drugs (NSAIDs), drug transport, human serum albumin

## Abstract

Ketoprofen is a popular non-steroidal anti-inflammatory drug (NSAID) transported in the bloodstream mainly by serum albumin (SA). Ketoprofen is known to have multiple side effects and interactions with hundreds of other drugs, which might be related to its vascular transport by SA. Our work reveals that ketoprofen binds to a different subset of drug binding sites on human SA than has been observed for other species, despite the conservation of drug sites between species. We discuss potential reasons for the observed differences in the drug’s preferences for particular sites, including ketoprofen binding determinants in mammalian SAs and the effect of fatty acids on drug binding. The presented results show that the SA drug sites to which a particular drug binds cannot be easily predicted based only on a complex of SA from another species and the conservation of drug sites between species.

## 1. Introduction

Serum albumin (SA) is the most abundant protein in mammalian blood plasma, with a concentration in human blood typically varying from 35 to 50 g/L (Doweiko & Nompleggi, 1991). The structure of SA consists of three homologous domains containing multiple pockets that can accommodate the binding of various classes of small molecules. Thanks to its high blood concentration and flexible structure, SA can serve as fatty acid, metal ion, and drug transporter in the blood (Peters, 1995; Lombardo *et al*, 2018). More than 600 FDA-approved drugs have been reported to bind to plasma proteins, mainly albumin, at levels of more than 50% of a total drug concentration bound (Lombardo *et al*, 2018). However, only about 30 FDA-approved drugs have structures of SA complexes available in the Protein Data Bank (PDB), including 25 drugs in complex with human SA. Ten binding sites within SA have been characterized as drug sites, and nine of these have been demonstrated to bind at least three FDA-approved drugs (Czub *et al*, 2020).

The binding of drugs to plasma proteins, mainly SA, is routinely evaluated during drug lead optimization (Bohnert & Gan, 2013; Trainor, 2007). Because of the high SA concentration in the blood, drug binding to SA is one of the significant factors determining their free plasma concentration. SA acts as a transporting reservoir for drugs, delivering drugs to the sites of action, extending their half-life in the blood, and contributing to their therapeutic efficiency by decreasing their aggregation. However, strong drug binding to SA may negatively affect its distribution in the body and is undesired in some cases. Only drug molecules that are not bound to plasma proteins are expected to be pharmacologically active (Bohnert & Gan, 2013; Trainor, 2007). Due to the high conservation of the sequence and structure of SA among species, its drug-binding properties are typically expected to be similar in different organisms. Nevertheless, a significant difference in binding to plasma proteins has been observed for some drugs for human and animal models (e.g., valproate in mouse, cefotetan in rats), which was linked to differences in binding to SA (Kosa *et al*, 1997; Acharya *et al*, 2006; Colclough *et al*, 2014).

Ketoprofen is a non-steroidal anti-inflammatory drug used to treat rheumatoid arthritis, osteoarthritis, dysmenorrhea, and alleviate moderate pain primarily in humans (Ketoprofen entry in Drugbank) and domesticated animals such as cats, dogs, and horses (Lees *et al*, 2003; Hazewinkel *et al*, 2003; Owens *et al*, 1995). Ketoprofen possesses a chiral center (Figure 1) and is typically administered orally or topically in the form of a racemic mixture. The (*S)*-enantiomer is primarily responsible for inhibiting prostaglandin synthesis, while the (*R)*-enantiomer is responsible for analgesic activity (Cooper *et al*, 1998; Ghezzi *et al*, 1998). However, it has been observed that some fraction of *(R)*-ketoprofen undergoes metabolic conversion to (*S)*-enantiomer in patients (Lorier *et al*, 2016). Ketoprofen is a commonly used over-the-counter NSAID, but multiple side-effects have been reported for this drug (Kantor, 1986; Loet, 1989). Moreover, ketoprofen is known to have 70 major and 260 moderate interactions with other drugs (Ketoprofen entry on Drugs.com). The side-effects of ketoprofen and its interactions with other medications may make it undesirable for some patients.

**Figure 1.**
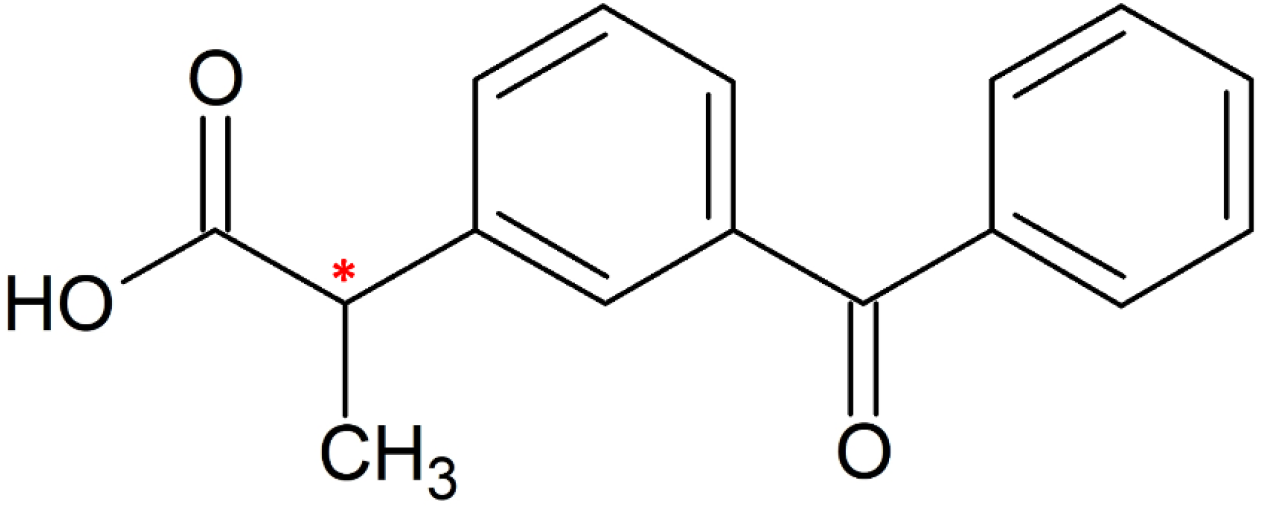
Structure of ketoprofen; the chiral center is labeled with an asterisk.

About 99% ketoprofen in the human blood is bound to SA (Ketoprofen entry in Drugbank), and its binding to mammalian SAs has been extensively studied using equilibrium dialysis (Dubois *et al*, 1993), calorimetry (Zielinski *et al*, 2020; Misra & Kishore, 2013), and spectroscopic methods (Maciążek-Jurczyk, 2014; Bi *et al*, 2011). These studies reported relatively similar ketoprofen binding affinities to human SA (Bi *et al*, 2011) (HSA; K_d_= 85 µM at 288.15 K), bovine SA (Misra & Kishore, 2013) (BSA; K_d1_=30 µM, K_d2_=189 µM), and leporine SA (Zielinski *et al*, 2020) (LSA; K_d_=49 µM). Recently, X-ray crystallography was employed to determine structures of ketoprofen complexes with BSA (Castagna *et al*, 2019) (PDB ID: 6QS9), ESA (Czub *et al*, 2020) (PDB ID: 6U4R), and LSA (Zielinski *et al*, 2020) (PDB ID: 6OCK). However, despite the high sequence and secondary structure conservation of albumin between species (Figure S1), none of the prior studies could provide conclusive information about the location of ketoprofen binding sites in human serum albumin (HSA). Accurate knowledge of the locations of ketoprofen binding sites on HSA would allow for a better understanding of its vascular transport, including the effects of non-enzymatic glycation caused by diabetes (Anguizola *et al*, 2013) and competition with other drugs for the same binding sites that may cause drug-drug displacement (Czub *et al*, 2020; Bohnert *et al*, 2010).

We present the first crystal structure of HSA in complex with ketoprofen and provide insights on the molecular basis of its transport in the blood. We also compare ketoprofen binding modes observed in other mammalian SAs and discuss potential reasons for the observed interspecies differences in its binding to SA.

## 2. Materials and methods

### 2.1 Materials

Recombinant HSA expressed in *Pichia pastoris* was purchased from Sigma-Aldrich (#A7736; ≥90% purity; St. Louis, MO, USA) as lyophilized powders and purified further as described below. According to the vendor, the construct has a single deletion of Asp from the N-terminus (Asp1) to create a hypoallergenic construct by eliminating the principal copper and nickel binding site of albumin. In addition, Cys34 was blocked by adding free cysteine to improve stability, monomer content, and homogeneity. Ketoprofen was purchased from Santa Cruz Biotechnology (#205359; ≥99% purity; Dallas, TX, USA) in the form of a racemic mixture, which is the same as the commercially available formulation of this drug.

### 2.2. Structure determination

#### 2.2.1. Protein purification for crystallization

HSA was dissolved in a buffer containing 50 mM Tris (pH 7.4) and 20 mM NaCl and subjected to gel filtration using the same buffer on a Superdex 200 column attached to an ÄKTA FPLC (GE Healthcare) at 4 °C. HSA concentration was estimated spectrophotometrically by measuring the absorbance at 280 nm with a Nanodrop 2000 (Thermo Scientific) using extinction coefficients ε_280-HSA_= 34,440 M^−1^cm^−1^ and molecular weight MW_HSA_= 66,470 kDa. Collected fractions of monomeric HSA were combined and concentrated to 162 mg/mL using an Amicon Ultra Centrifugal Filter (Millipore Sigma, #UFC903024) with a 30K MWCO.

#### 2.2.2. Protein crystallization

Crystallization was performed in 96-well plates (Hampton Research, HR3-123) that were set up using a Mosquito crystallization robot (TTP Labtech). Prior to crystallization, HSA solution at concentration of 162 mg/mL (dissolved in 50 mM Tris pH 7.4 and 20 mM NaCl) was mixed with 100 mM ketoprofen in 100% DMSO in ratio 9:1 (final ketoprofen concentration 10 mM) and incubated for several hours at 37 °C. Aliquots of 0.2 µL of the resulting HSA-ketoprofen solution were mixed with 0.2 µL aliquots of reservoir solution (24% (w/v) PEG 3350, 50 mM potassium phosphate at pH 7.0). The crystallization plate was incubated at room temperature for 3 months and then at 37 °C for several days until the first crystals were observed. Harvested crystals were flash-cooled without any additional cryoprotectant.

#### 2.2.3. Data collection and structure determination

Data collection was performed from a single crystal at the SBC 19-ID beamline at the Advanced Photon Source, Argonne National Laboratory (Argonne, IL). The experiment was performed at 100 K, using X-rays with wavelength 0.979 Å. HKL-3000 (Minor *et al*, 2006; Otwinowski & Minor, 1997) was used to process, integrate, and scale the data. Corrections for radiation decay and anisotropic diffraction were applied (Borek *et al*, 2010). The native structure of HSA (PDB ID: 4K2C) was used as the template for molecular replacement. Structure determination and refinement were performed using HKL-3000 integrated with MOLREP (Vagin & Teplyakov, 2010), REFMAC (Murshudov *et al*, 2011), Coot (Emsley *et al*, 2010), and other programs from the CCP4 package (Winn *et al*, 2011). The refinement process followed recent state-of-the-art guidelines (Shabalin *et al*, 2018; Majorek *et al*, 2020). Thirteen TLS groups, determined by the TLS Motion Determination Server, were applied during refinement (Painter & Merritt, 2006). (*S*)-or (*R*)-enantiomers of ketoprofen were chosen by careful evaluation of each candidate fit to the 2mFo-DF, and mFo-DFc omit maps (calculated for 10 cycles of REFMAC refinement without the ligand). Each choice was supported by comparing fit to the after-refinement maps, the resulting ADP values, and the interactions with the protein (hydrogen bonds, salt bridges, and the lack of clashes). Partial occupancy was evaluated for the (*R*)-ketoprofen molecule in drug site 9, which resulted in the appearance of positive electron density and comparatively low ADP values; therefore, the occupancy was kept at 100%. The ACHESYM server (Kowiel *et al*, 2014) was used for the standardized placement of the model in the unit cell. The PISA server (Krissinel & Henrick, 2007) was used to analyze the residues involved in interactions between the ligand and macromolecule. PyMOL (The PyMOL Molecular Graphics System, Version 1.5.0.3 Schrödinger, LLC) and ChemSketch were used for figure generation. The Dali server (Holm, 2019) was used for the structure comparison and calculation of Cα RMSD. The statistics for diffraction data collection, structure refinement, and structure quality are summarized in **Table 1**. Diffraction images are available at the Integrated Resource for Reproducibility in Macromolecular Crystallography at http://proteindiffraction.org (Grabowski *et al*, 2016) with DOI: 10.18430/m37jwn. The atomic coordinates and structure factors were deposited in the PDB with accession code 7JWN.

**Table 1.** Data collection, structure refinement, and structure quality statistics.

|  |  |
| --- | --- |
| **PDB ID** | 7JWN |
| **Diffraction images DOI** | 10.18430/m37jwn |
| **Resolution (Å)** | 50.00-2.60<br>(2.64-2.60) |
| **Wavelength (Å)** | 0.979 |
| **Space group** | C2 |
| **Unit-cell dimensions: a, b, c (Å)** | 170.5, 38.9, 98.5 |
| **Angles: $\alpha$, $\beta$, $\gamma$ (°)** | 90.0, 104.5, 90.0 |
| **Protein chains in the ASU** | 1 |
| **Completeness (%)** | 96.4 (88.5) |
| **Number of unique reflections** | 18925 (851) |
| **Redundancy** | 4.2 (3.5) |
| **$\langle I \rangle / \langle \sigma(I) \rangle$** | 16.9 (1.3) |
| **CC $\frac{1}{2}$** | (0.60) |
| **R<sub>merge</sub>** | 0.081 (0.803) |
| **R<sub>meas</sub>** | 0.093 (0.925) |
| **R<sub>work</sub>/R<sub>free</sub>** | 0.183 / 0.231 |
| **Bond lengths rmsd (Å)** | 0.002 |
| **Bond angles rmsd (°)** | 1.1 |
| **Mean ADP (Å<sup>2</sup>)** | 52 |
| **Mean ADP for ketoprofen molecules (Å<sup>2</sup>)** | (S)-ketoprofen: 24.4 (DS2),<br>16.6 (DS3; sub-site A),<br>47.1 (DS3; sub-site B);<br>(R)-ketoprofen: 72.1 (DS9) |
| **Number of protein atoms** | 4646 |
| **Mean ADP for protein (Å<sup>2</sup>)** | 53 |
| **Number of water molecules** | 192 |
| **Mean ADP for water molecules (Å<sup>2</sup>)** | 36 |
| **Clashscore** | 1.27 |
| **MolProbity score** | 1.07 |
| **Rotamer outliers (%)** | 0.59 |
| **Ramachandran outliers (%)** | 0.0 |
| **Ramachandran favored (%)** | 96.23 |

Values in parentheses are for the highest resolution shell. Ramachandran plot statistics are calculated by MolProbity (Williams *et al*, 2018). DS2, DS3, and DS9 refer to drug-binding sites 2, 3, and 9.

## 3. Results

The crystal of the HSA complex with ketoprofen grew in the *C*2 space group and contained one protein chain in the asymmetric unit. The protein model is complete except for the first residue (Ala2), for which the electron density is not observed. The electron density revealed binding of one (*S*)-ketoprofen molecule to drug site 2, two (*S*)-ketoprofen molecules to drug site 3, and one (*R*)-ketoprofen molecule to drug site 9 (Figure 2). All three sites were previously reported to bind multiple FDA-approved drugs (Figure S2) (Czub *et al*, 2020). The structure also contains three molecules of fatty acids, modeled as myristic acid, bound to FA3 (which overlaps with drug site 2), drug site 5 (not previously characterized as a fatty acid binding site), and FA5 (overlaps with drug site 8). Accordingly, the determined HSA complex with ketoprofen has an almost identical conformation to HSA complexed with myristic acid (PDB ID: 1BJ5). However, it is noticeably different from a ligand-free HSA structure (PDB ID: 4K2C), as can be concluded from the RMSD values between the aligned Cα atoms (Table S1, Figure S3). Fatty acids were not added during crystallization and are most likely remnants from the purification. Free cysteine was added by the manufacturer to the protein during purification to block the sole free cysteine residue in HSA and prevent albumin dimerization. Based on the observed electron density, Cys34 forms a disulfide bond with another molecule of cysteine. The quality of electron density observed for ligands in the determined structure can be inspected interactively at https://molstack.bioreproducibility.org/project/view/VW8s7hb1Z9mnCLbg3NBU/. As a control, we also determined a 2.70 Å structure of HSA obtained from the same crystallization conditions but not containing ketoprofen (*P*1 space group; unit cell dimensions: 38.0 Å, 86.2 Å, 97.3 Å; angles: 75.0**°**, 89.6**°**, 78.6**°**; data not shown). In the control structure, all ketoprofen binding sites (drug sites 2, 3, and 9) remain unoccupied, Cys34 also forms a disulfide bond with another molecule of cysteine, and fatty acids bind to the same sites as in the HSA-ketoprofen complex.

**Figure 2.**
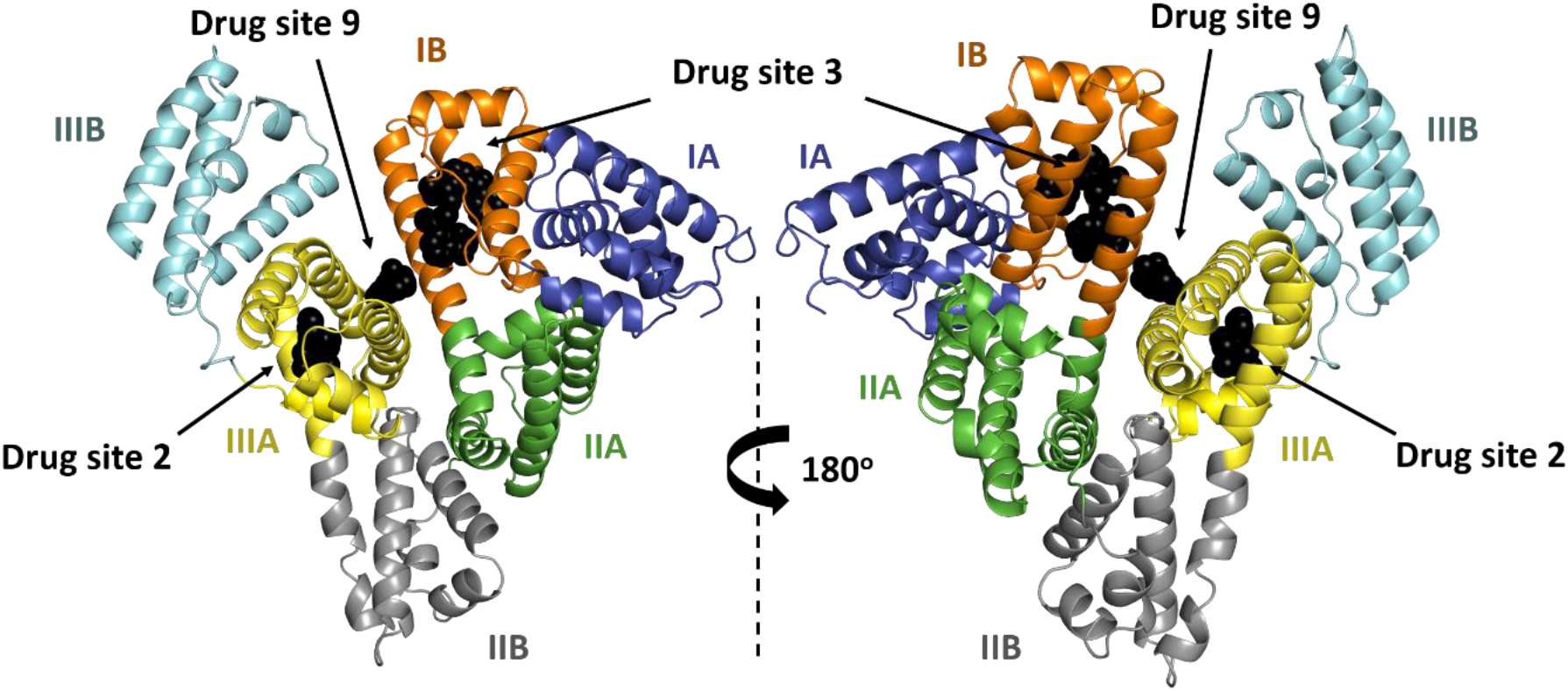
The overall structure of HSA complex with ketoprofen. Albumin subdomains are each shown in a different color. Roman numerals (I, II, III) are associated with domains and letters (e. g., IB) with subdomains. Ketoprofen molecules are shown with atoms in black spheres.

### 3.1. Ketoprofen binding sites in HSA

Drug site 2, also known as Sudlow site II and FA3/FA4, is one of the three major drug binding sites in SA (Figure 3) (Czub *et al*, 2020; Sudlow *et al*, 1976, 1975). (*S*)-ketoprofen molecule at this site is stabilized by strong hydrophobic interactions with surrounding residues (mainly Tyr411, Val415, Val418, Leu423, Val426, Leu430, Leu453, Val456, Leu457, Leu460, Phe488) and also by hydrogen bonds between its carboxylate group and hydroxyl groups of Tyr411 and Ser489. Arg410 may also contribute a remote charge-charge interaction with the carboxylate group. All of the residues involved in binding of (*S*)-ketoprofen to HSA at drug site 2 are listed in Table An (*S*)-ketoprofen molecule bound to drug site 2 overlaps with the fatty acid previously reported to bind in FA4 (see PDB ID: 1BJ5) (Curry *et al*, 1998), and is located close to FA3, which is occupied by a molecule of myristic acid in the reported structure.

**Figure 3.**
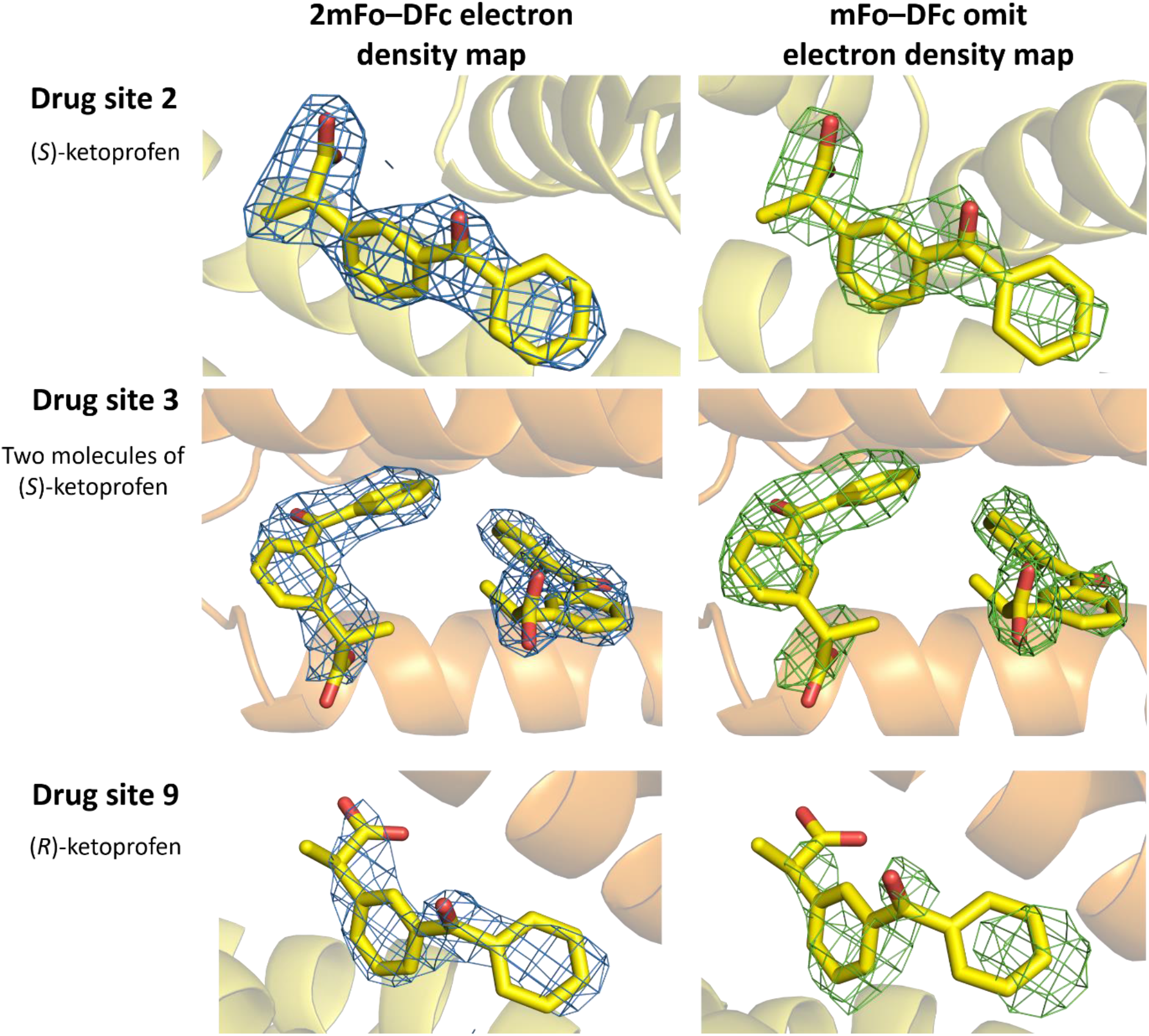
Ketoprofen binding sites in human serum albumin (PDB ID: 7JWN) with 2mFo–DFc electron density map (RMSD 1.0) presented in blue and mFo–DFc omit electron density map (map calculated after 10 REFMAC refinement cycles without the drug in the model, RMSD 2.5) presented in green. Ketoprofen molecules are shown in stick representation with oxygen atoms in red and carbon atoms in yellow. The colors of the helices correspond to the colors in Figure 2. The electron density and the model can be inspected interactively at https://molstack.bioreproducibility.org/project/view/VW8s7hb1Z9mnCLbg3NBU/.

Drug site 3, also called the oncological drug site and FA1 (Wang *et al*, 2013), is also one of the three major drug binding sites on SA (Czub *et al*, 2020; Sudlow *et al*, 1975, 1976). This site has two (*S*)-ketoprofen molecules bound at the same time, and due to that, can be divided into two sub-sites. Sub-site A overlaps with previously characterized FA1 (Curry *et al*, 1998) and has (*S*)-ketoprofen bound. (*S*)-ketoprofen at sub-site A is stabilized by strong hydrophobic interactions with residues forming a narrow binding pocket (mainly Leu115, Met123, Phe134, Tyr138, Leu139, Ile142, Leu154, Ala158, Tyr161, Phe165, and Leu182), by a salt bridge between its carboxylate group and Arg117’s guanidino group, and a hydrogen bond of the carboxylate group with Tyr161’s hydroxyl group (Table 2). Moreover, a remote charge-charge interaction of the carboxylate group with Arg186 is likely an additional stabilizing factor. Sub-site B within drug site 3 harbors an (*S*)-ketoprofen molecule surrounded by sparse hydrophobic residues, mainly Ile142, Phe149, Leu154, Phe157, Tyr161, and aliphatic part of Lys190’s side chain. At this sub-site, (*S*)-ketoprofen’s carboxylate group forms a hydrogen bond with the His146 side chain (NE2 atom) and a remote charge-charge interaction with Arg145. As compared to drug site 2 and the sub-site A, the sub-site B offers significantly smaller hydrophobicity (as can be seen by the significantly lower number of hydrophobic residues taking part in the interaction) and weaker hydrophilic interactions (no salt bridges and only one hydrogen bond), which may suggest weaker binding of (*S*)-ketoprofen. Indeed, the high atomic ADP values observed for this ligand (Table 1) may suggest its partial occupancy but may also be a result of its positional variability between HSA molecules in the crystal. Notably, the molecules of (*S*)-ketoprofen bound to the sub-sites of drug site 3 have their phenyl rings located within 4 Å of each other (Figure 3), suggesting that this hydrophobic interaction additionally stabilizes both (*S*)-ketoprofen molecules and may result in possible cooperativity in binding.

**Table 2.**
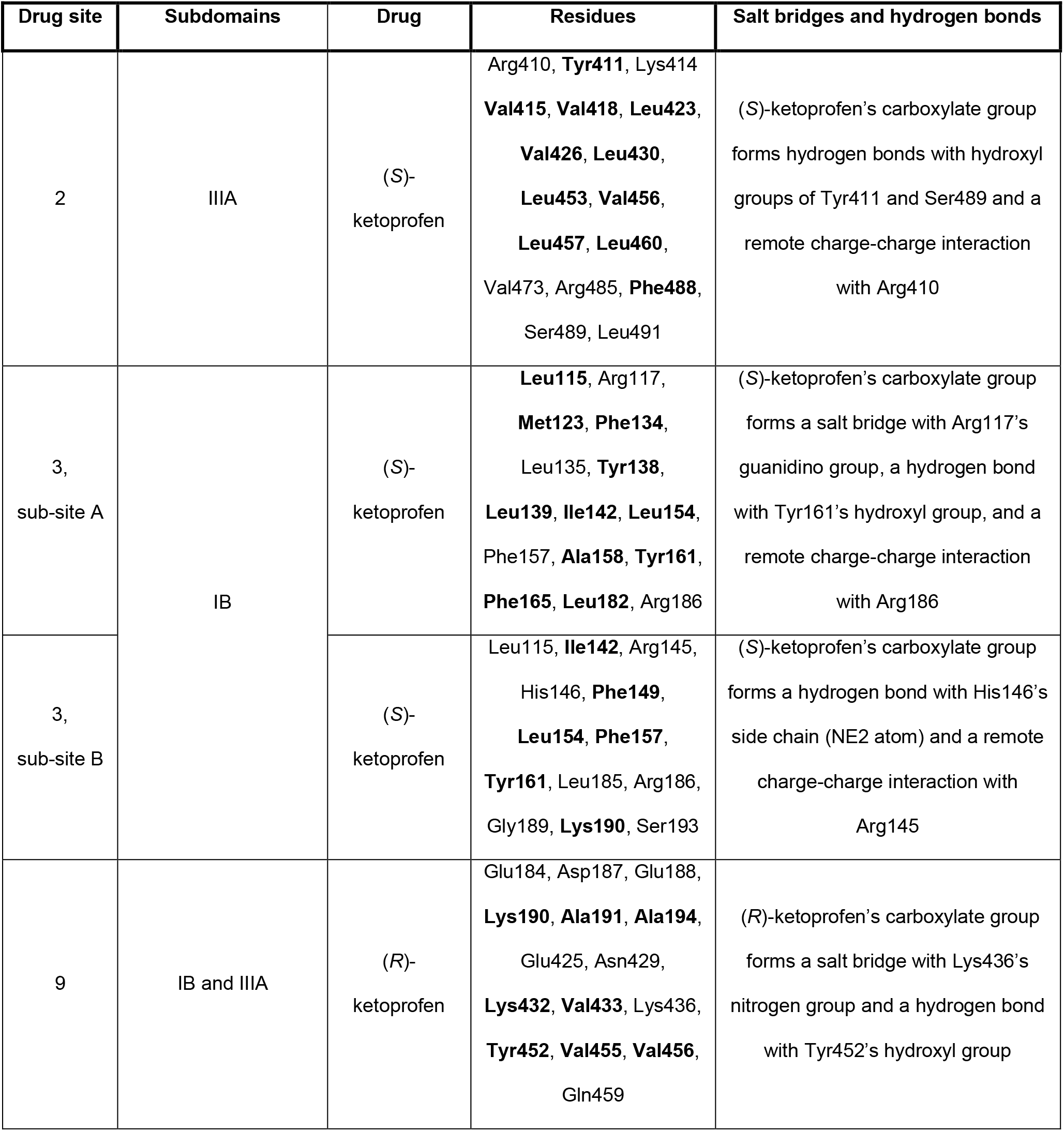
The residues that participate in binding of ketoprofen to human serum albumin and hydrophilic interactions observed in ketoprofen binding sites. Residues shown in bold are responsible for a major hydrophobic contribution to drug binding.

Drug site 9, located near FA8 and FA9, is a much less common drug binding site on SA (Czub *et al*, 2020). This site has the only (*R*)-ketoprofen molecule in the reported structure. The (*R*)-ketoprofen molecule is stabilized by some hydrophobic interactions (mainly Ala191, Ala194, Val433, Tyr452, Val455, Val456, and aliphatic parts of Lys190 and Lys432), a hydrogen bond formed by its carboxylate group with Tyr452’s hydroxyl group, and a salt bridge between the carboxylate group and Lys436’s nitrogen group. The (*R*)-ketoprofen molecule has relatively high ADP values, suggesting partial occupancy or positional variability, which is also correlated with a significantly smaller hydrophobicity of this site that likely results in a lower affinity of this site.

### 3.2. Comparison of ketoprofen binding sites in HSA and other mammalian SAs

The structure of the HSA complex with ketoprofen and the observed binding sites were compared to previously reported structures of ESA, BSA, and LSA complexes with ketoprofen. The comparison of the experimental conditions that were used and the contents, if any, of the known albumin drug sites is shown in Table 3.

**Table 3.** Conditions used for crystallization of SA-ketoprofen complexes and the ligands observed in the common drug binding sites. N/R – not reported.

|  | **HSA-ketoprofen**<br>(PDB ID: 7JWN) | **ESA-ketoprofen** (Czub<br>*et al*, 2020)<br>(PDB ID: 6U4R) | **BSA-ketoprofen**<br>(Castagna *et al*,<br>2019)<br>(PDB ID: 6QS9) | **LSA-ketoprofen**<br>(Zielinski *et al*,<br>2020)<br>(PDB ID: 6OCK) |
| --- | --- | --- | --- | --- |
| **SA source** | Recombinant HSA expressed in *Pichia pastoris* (Sigma # A7736) | ESA isolated from horse blood (Equitech-Bio #ESA62) | BSA isolated from bovine blood (Sigma # N/R) | LSA isolated from leporine blood (Sigma # N/R) and defatted prior to the experiment |
| **Crystallization drops** | 162 mg/mL HSA solution in a buffer containing 50 mM Tris (pH 7.5) and 20 mM NaCl was mixed with 100 mM ketoprofen in DMSO in ratio 9:1. Aliquots of 0.2 μL of the HSA solution were mixed with 0.2 μL aliquots of reservoir solution. | 1 μL of 34 mg/mL ESA solution in a buffer containing 10 mM Tris (pH 7.5) and 150 mM NaCl was mixed with 1 μL aliquots of reservoir solution. ESA crystals were soaked with ketoprofen suspended in DMSO. | 10 mg/mL BSA solution in a buffer containing 10 mM Tris (pH 7.5) and 150 mM NaCl was mixed with ketoprofen suspended in ethanol. Then, the BSA solution was mixed with the reservoir solution. | 67 mg/mL LSA solution in a buffer containing 10 mM Tris (pH 7.4) and 100 mM NaCl was mixed with ketoprofen suspended in ethanol. Then, the LSA solution was mixed with the reservoir solution. |
| **Final ketoprofen concentration** | ~5.0 mM | ~3.0 mM | ~0.7 mM | ~4.6 mM |
| **Reservoir solution** | 24% PEG 3350, 50 mM potassium phosphate buffer at pH 7.0 | 0.2 M lithium sulfate, 2.0 M ammonium sulfate, 0.1 M Tris buffer at pH 7.4 | 18% PEG MME 5000, 0.2M ammonium chloride, 0.1M MES buffer at pH 6.5 | 8% polypropylene glycol 400, 16% PEG 3350, 0.2 M ammonium acetate, and 0.1 M Tris buffer at pH 8.0 |
| **DS1** | - | UNL | (*R*)-ketoprofen | Acetate ion |
| **DS2** | (*S*)-ketoprofen; fatty acid | Fatty acid | - | (*S*)-ketoprofen |
| **DS3** | Two molecules of (*S*)-ketoprofen | - | - | PEG molecule |
| **DS4** | - | (*S*)-ketoprofen | - | Polymer with PDB code 2J3 |
| **DS5** | Fatty acid | - | - | - |
| **DS6** | - | (*S*)-ketoprofen | - | (*S*)-ketoprofen, acetate ion |
| **DS7** | - | - | - | Polymer with PDB code POG |
| **DS8** | Fatty acid | - | - | - |
| **DS9** | (*R*)-ketoprofen | - | - | Acetate ion |
| **DS10** | - | (*S*)-ketoprofen | - | - |

#### 3.2.1. Ketoprofen binding sites in ESA

ESA (the pairwise sequence identity to HSA is 76.1%) has recently been reported to bind (*S*)-ketoprofen at drug sites 4, 6, and 10 (Figure 4) (Czub *et al*, 2020). Surprisingly, these drug sites are unoccupied in the HSA structure reported herein. The conservation of residues comprising these drug sites in ESA and HSA has been discussed in detail by Czub et al (Czub *et al*, 2020), which concluded that drug site 4 significantly differs between ESA and HSA (57% conservation), drug site 6 is partially conserved (75% of residues is conserved), and drug site 10 is very well conserved between albumin from both species (94%). Therefore, the lack of (*S*)-ketoprofen in sites 4 and 6 may be attributed to these differences. However, all the residues involved in binding of (*S*)-ketoprofen in ESA at drug site 10 are the same in HSA, except for Ile7, which is modified to Val. Moreover, Ile7 in ESA contributes only to hydrophobic interactions with the drug molecule, further suggesting conservation of the site and leading to the expectation that drug site 10 in HSA may bind (*S*)-ketoprofen as well, albeit it was not observed in the structure reported herein. Drug site 2, where (*S*)-ketoprofen binds to HSA, is occupied by the molecule of fatty acid in the structure of the ESA-ketoprofen complex (PDB ID: 6U4R) and potentially prevent drug binding. Drug sites 3 and 9 remain unoccupied in the ESA-ketoprofen structure (PDB ID: 6U4R).

**Figure 4.**
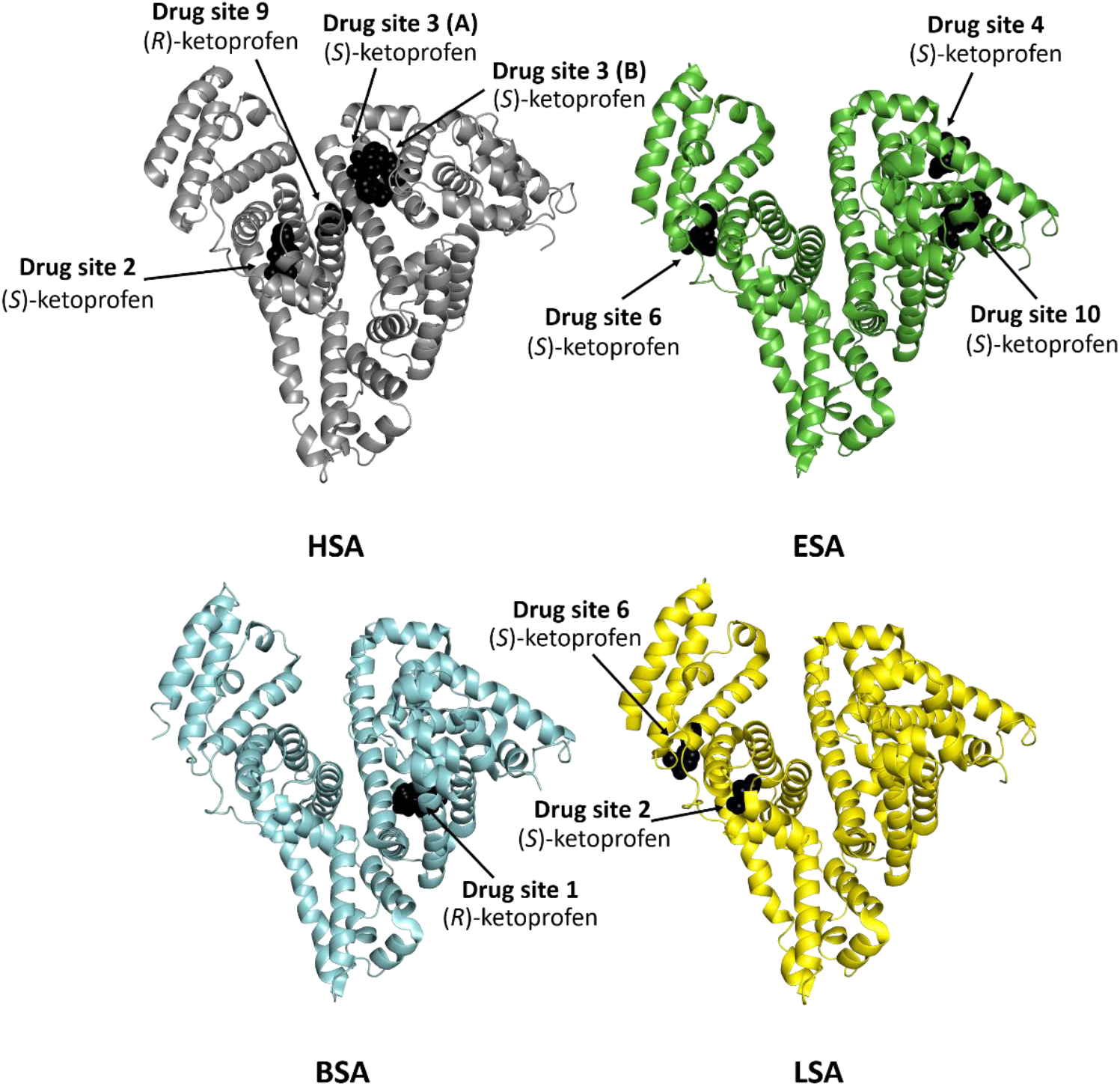
Ketoprofen binding sites in mammalian serum albumins. Structures of ketoprofen complexes with HSA (PDB ID: 7JWN), ESA (PDB ID: 6U4R), BSA (PDB ID: 6QS9), LSA (PDB ID: 6OCK).

#### 3.2.2. Ketoprofen binding sites in BSA

Drug site 1 (Sudlow site I) has been reported as the only ketoprofen binding site in BSA (the pairwise sequence identity to HSA is 75.6%), and the drug molecule was modeled as (*R*)-enantiomer at this site (PDB ID: 6QS9, Figure 3) (Castagna *et al*, 2019). Most of the residues involved in interactions with (*R*)-ketoprofen at drug site 1 are conserved between BSA and HSA (Figure 5), including Arg256 (Arg257 in HSA) and Tyr149 (Tyr150 in HSA). These residues form a salt bridge and a hydrogen bond with ketoprofen’s carboxylate group, respectively. Only two residues are different, Arg198 (Lys199 in HSA) and Lys221 (Arg222). Moreover, they are involved only in hydrophobic interactions, and these changes should not affect the binding of ketoprofen to this site. However, despite the very high sequential conservation of drug site 1 between BSA and HSA (89%), this site remains unoccupied in the structure of the HSA-ketoprofen complex (PDB ID: 7JWN). Drug sites 2, 3, and 9 are free of ligands in the BSA-ketoprofen structure (PDB ID: 6QS9). Binding of only one ketoprofen molecule to BSA may be explained by a lower ketoprofen concentration than those used for other complexes (Table 3).

**Figure 5.**
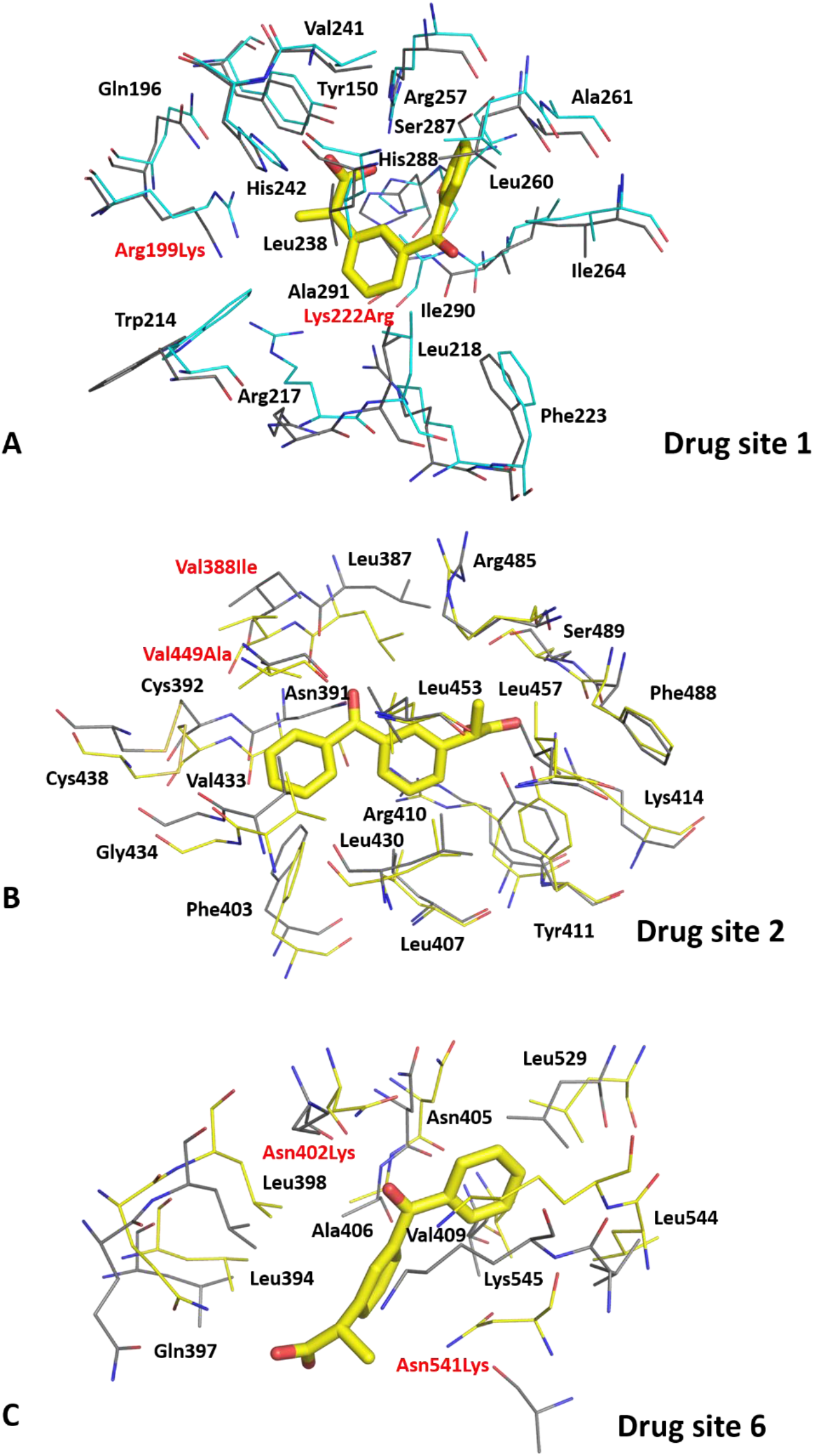
Superposition of ketoprofen binding sites in BSA (panel A, PDB ID: 6QS9) and LSA (panels B and C, PDB ID: 6OCK) with analogous sites in ligand-free HSA (PDB ID: 4K2C). Carbon atoms in BSA, LSA, and HSA are shown in cyan, yellow, and gray, respectively. Residue numbers correspond to positions in HSA. Residues written in black are conserved between BSA or LSA and HSA, while those written in red differ. The naming scheme for differing residues is as follows: residue from BSA or LSA, residue number, residue from HSA.

#### 3.2.3. Ketoprofen binding sites in LSA

LSA (the pairwise sequence identity to HSA is 73.4%) has been reported to bind (*S*)-ketoprofen in drug sites 2 and 6 (PDB ID: 6OCK, Figure 4) (Zielinski *et al*, 2020). Surprisingly, drug site 2 binds (*S*)-ketoprofen in both HSA and LSA, but the binding mode is different (Figure 6). The (*S*)-ketoprofen molecule at drug site 2 in LSA is stabilized by hydrophobic interactions with surrounding residues and forms a salt bridge and a hydrogen bond between its carboxylate group and Lys414, and Tyr411, respectively. Most of the residues involved in these interactions are conserved (89%). Two residues that differ between LSA and HSA (Val388 is replaced by Ile and Val449 by Ala) do not change the character of the binding site (Figure 5). The drug’s carboxylate group occupies roughly the same position, but its hydrophobic parts are oriented in opposite directions. Most likely, the presence of a fatty acid molecule at this site (FA3) in the HSA structure reported herein affects the conformation of (*S*)-ketoprofen. Previously, (*S*)-ibuprofen has been reported to bind to drug site 2 in HSA and ESA with two different binding modes that resemble those of (*S*)-ketoprofen (Figure S4) (Czub *et al*, 2020).

**Figure 6.**
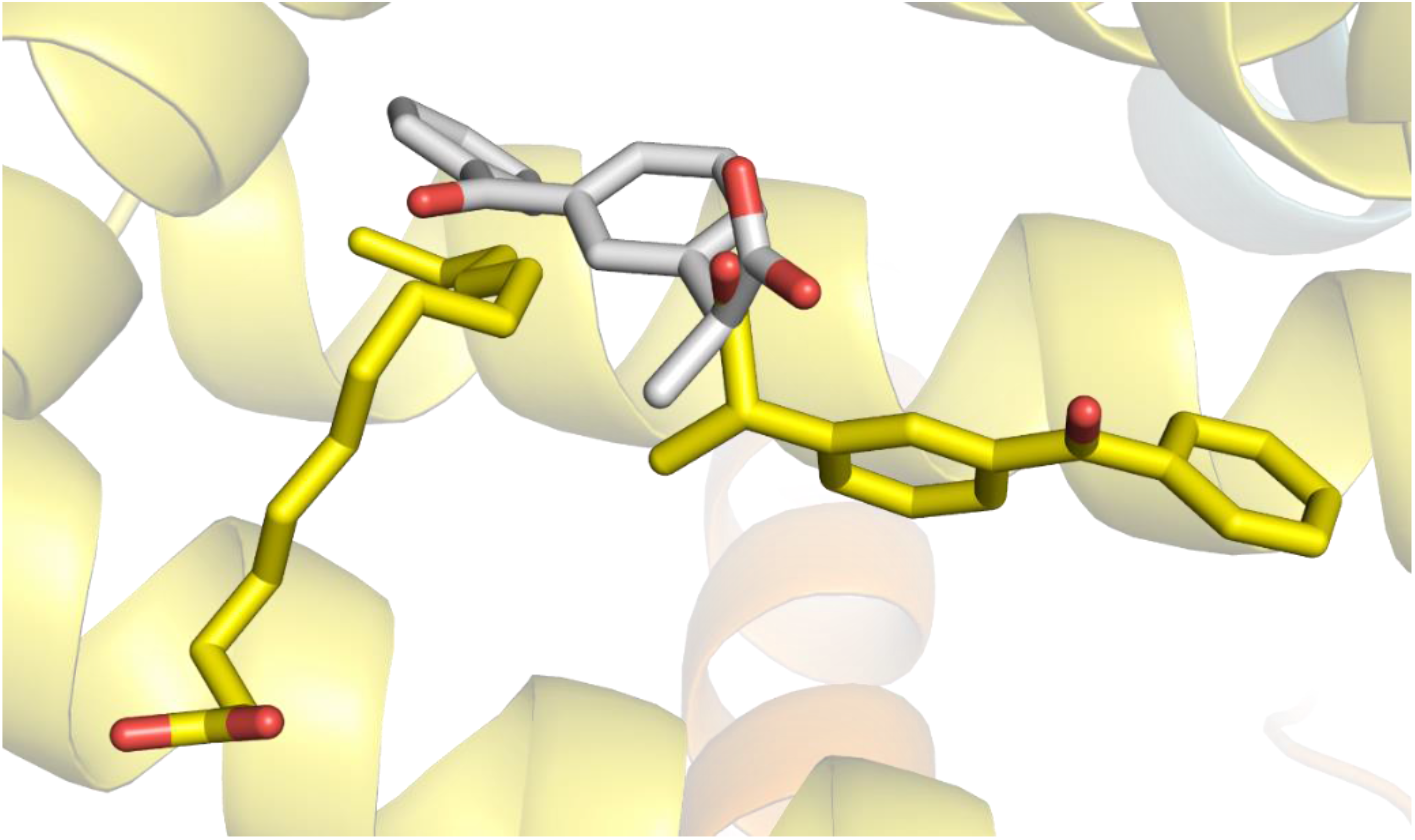
Comparison of (*S*)-ketoprofen binding to drug site 2 in HSA (PDB ID: 7JWN) and LSA (PDB ID: 6OCK). (*S*)-Ketoprofen molecule and molecule of a fatty acid bound to HSA are shown in stick representation with oxygen atoms in red and carbon atoms in yellow, while a molecule of (*S*)-ketoprofen bound to LSA is shown in stick representation with oxygen atoms in red and carbon atoms in gray. The colors of the helices correspond to the colors in Figure 2.

The other molecule of (*S*)-ketoprofen binds to drug site 6 in LSA, where it is stabilized by hydrophobic interactions and by hydrogen bonds between its carboxylate group and Asn397’s sidechain (NE2 atom) and between its carbonyl group and sidechains of Asn402 (ND2 atom) and Lys545. This binding site is 82% conserved between LSA and HSA. Two residues that are modified (Asn402 to Lys and Asn541 to Lys) change the overall charge in the cavity, and due to the bigger size of the side chain may affect the drug’s conformation or even prevent its binding. Drug site 6 is unoccupied in the HSA-ketoprofen structure (PDB ID: 7JWN). In the LSA-ketoprofen structure (PDB ID: 6OCK), drug site 3 is occupied by a molecule of a polyethylene glycol, which may prevent drug binding to this site, and an acetate ion is present in drug site 9.

## 4. Discussion

We have determined the first structure of the ketoprofen complex with HSA and revealed that four ketoprofen molecules bind to drug sites 2, 3, and 9 therein. The electron density observed for ketoprofen molecules in the determined structure indicated binding of (*S*)-enantiomers at drug sites 2 and 3 (two molecules) and an (*R*)-enantiomer at drug site 9. We compared the determined structure to previously reported ketoprofen complexes with SA from other mammals, where ketoprofen was shown to bind to drug site 4, 6, and 10 in ESA (PDB ID: 6U4R) (Czub *et al*, 2020), drug site 1 in BSA (PDB ID: 6QS9) (Castagna *et al*, 2019), and drug sites 2 and 6 in LSA (PDB ID: 6OCK; Figure 4) (Zielinski *et al*, 2020). Despite high sequence identity (the pairwise sequence identity to HSA is 76.1% for ESA, 75.6% for BSA, and 73.4% for LSA; Figure S1), well-conserved binding sites (Figure 5), and similar ketoprofen binding affinity (Bi *et al*, 2011; Zielinski *et al*, 2020; Misra & Kishore, 2013), drug site 2 is the only site that was observed to bind ketoprofen in HSA and a SA from another species (LSA). Residues involved in ketoprofen binding to drug site 2 are 89% conserved between LSA and HSA, and only two of them differ (in LSA, Ile388 is replaced by Val and Ala449 by Val). These modified residues contribute to hydrophobic interactions and their alterations do not change the character of this site significantly. However, the structures discussed herein show that the SA drug sites to which a particular drug is binding cannot be easily predicted based only on a known complex of SA from another species and the conservation of drug sites between species. For instance, drug site 10 is very well conserved between ESA and HSA (94% of residues is conserved; only Ile7 is modified to Val), including the character and protein fold, but ketoprofen binding to this site was observed only for ESA. Therefore, there are additional factors that define which drug sites we see occupied in the discussed structures, not just sequence conservation.

These variations in the binding sites where ketoprofen binds in different structures can be at least partially explained by different crystallization conditions (Table 3) used to produce these structures. In the structure of LSA (PDB ID: 6OCK), some of the biding sites are occupied by compounds used as a part of crystallization conditions (e.g., drug site 3 is occupied by a molecule of polyethylene glycol). Moreover, the ESA-ketoprofen complex is the only complex that was crystallized from a high-salt condition without PEGs present. All other complexes discussed herein were obtained from conditions with 16-24% of PEG and with much lower salt concentrations. Perhaps, these and other differences in conditions (e.g., the presence of different organic solvents used to dissolve the drugs) have resulted in the difference in occupied sites observed, as ionic strength, viscosity, and temperature have been shown to affect ligand binding to proteins (Papaneophytou *et al*, 2014).

Another potential reason for these variations can be the variability in the presence of fatty acids and other compounds that are remnants from the purification. In the structures of HSA and ESA (PDB IDs: 7JWN and 6U4R, respectively), some drug sites are occupied by molecules of fatty acids that were not added during crystallization (see Table 3). For instance, drug site 2 in the ESA-ketoprofen complex (PDB ID: 6U4R) is occupied by the molecule of a fatty acid that potentially prevents drug binding. In addition to the direct competition of fatty acids for the drug binding sites, fatty acids also change the overall conformation of albumin, and affect the affinity of drugs for the binding sites (Curry *et al*, 1998; Petitpas *et al*, 2003). For example, HSA in the determined structure has a conformation significantly different from HSA without bound fatty acids and different from other SA complexes with ketoprofen (Figure S3). Calculated RMSD values are higher in the structural comparison of SA complexes with fatty acids and ligand-free SAs than in the comparison of SA complexes with ketoprofen and ligand-free SAs (Table S1). These results indicate that the presence of fatty acids alters the conformation of SA more significantly than the binding of ketoprofen. Similarly, other metabolites that interact with SA in physiological conditions (e.g., glucose) can occupy drug sites and even chemically modify SA’s structure, as in the case of non-enzymatic glycosylation. Due to that, SA isolated from natural sources may have different impurities present or different chemical modifications than SA expressed in yeasts. These modifications likely affect a small percentage of SA molecules and, as a result, are not usually observed in crystal structures (with the exception of fatty acids). However, the resulting heterogeneity may have an impact on the resulting crystal structure.

The concentration of drugs used for crystallization or crystal soaking can also be an important factor that affects the number of binding sites observed for a particular drug. It is known that SA’s drug sites have different affinities for ligands, and ketoprofen could bind to lower-affinity sites on SA at a sufficiently high occupancy to be seen on the electron density maps under a higher concentration of the drug. For instance, BSA has been observed to bind only one molecule of ketoprofen, whereas all other reported SA structures bind 2, 3, or even 4 ketoprofen molecules. If a higher concentration of ketoprofen was used in the BSA crystallization experiment, we would likely see other sites occupied by the drug. Also, we observed the binding of two ketoprofen molecules to drugs site 3 in HSA, but we do not know if it would be possible to observe the binding of only one drug molecule at this site if we had a lower concentration of ketoprofen due to a potential cooperativity effect. Moreover, the electron density observed for ketoprofen at drug site 9 is weaker than that observed at drug sites 2 and 3, which may be related to the lower binding affinity of the drug to this site, and potentially could disappear at a lower concentration. Additionally, the way the ligand was added to the protein, namely co-crystallization or crystal soaking, may affect where it binds to SA due to potentially different accessibility of sites in the solution vs. the crystal form. Together, this knowledge and observations suggest that SA may utilize different binding sites in crystal structures depending on experimental conditions and sites’ availability.

HSA is known to bind chiral drugs stereoselectively (Shen *et al*, 2013). In the reported structure of HSA, we observed the binding of three (*S*)-ketoprofen molecules (drug sites 2 and 3) and only one (*R*)-ketoprofen molecule (drug site 9). The shape of drug sites 2, 3, and 9 clearly supports the binding of the specific enantiomer of ketoprofen, suggesting differences in binding affinities of HSA to ketoprofen enantiomers. This observation agrees with the previous reports stating that HSA can bind (*R*)- and (*S*)-enantiomers of ketoprofen with different affinities, depending on the experimental conditions (Dubois *et al*, 1993; Zhivkova & Russeva, 1998). We can also conclude that SA typically promotes binding of (*S*)-profens, which was previously proposed (Czub *et al*, 2020; Zielinski *et al*, 2020), but its stereoselectivity is ultimately dictated by the structure of a drug and its fit to a particular binding site. The ten binding sites available on SA have a different affinity to different stereoisomers, and some of them may go against the general trend, as is in the case of site 9, which binds (*R*)-ketoprofen.

The results reported herein provide insights into the molecular basis of ketoprofen transport across species but also indicate a need for similar studies performed for other animal models. Determination of drug complexes with both HSA and SA from animals often used as model organisms would allow for the evaluation of these models and contribute to our understanding of differences in drug plasma binding in species. Our study also highlights the importance of multiple experimental conditions, such as the composition of the crystallization solutions, source of the protein, and the presence of fatty acids, that may affect protein’s conformation, availability/accessibility of binding sites and ultimately change what sites are occupied by a particular drug in the crystal structure. Generally, in structural biology, such differences are not expected to affect the binding of the specific binders to their sites. However, SA has not evolved to bind drugs specifically, which might allow for changes in drug preferences for a specific site depending on the conditions used and the availability of binding sites. Control for these variables and determining structures of complexes in varying conditions and with various SA sources is necessary for the reproducibility of albumin research.

## Contributions

M.P.C. determined the structure of the HSA-ketoprofen complex; M.P.C. and I.G.S. analyzed the obtained structure; M.P.C. and I.G.S., drafted the manuscript; M.P.C. prepared figures; M.P.C., I.G.S., and W.M. critically read, discussed, and edited the manuscript, and W.M. supervised and coordinated the entire project.

## Acknowledgments

We thank David R. Cooper for the critical reading and discussions of the manuscript. This work was supported by the National Institute of General Medical Sciences grants R01-GM132595, and federal funds from the National Institute of Allergy and Infectious Diseases, National Institutes of Health, United States Department of Health and Human Services under contract nos. HHSN272201200026C and HHSN272201700060C. M.P.C. acknowledges the support of the Robert R. Wagner Fellowship at the University of Virginia. We also thank Changsoo Chang for his assistance in data collection. Results shown in this report are derived from work performed at Argonne National Laboratory (ANL), Structural Biology Center (SBC) at the Advanced Photon Source (APS), under U.S. Department of Energy, Office of Biological and Environmental Research contract DE-AC02-06CH11357. Part of the results reported herein has been previously presented in the PhD thesis of M. P. C.

## References

Acharya MR, Sparreboom A, Sausville EA, Conley BA, Doroshow JH, Venitz J & Figg WD (2006) Interspecies differences in plasma protein binding of MS-275, a novel histone deacetylase inhibitor. Cancer Chemother Pharmacol 57: 275–81

Anguizola J, Matsuda R, Barnaby OS, Hoy KS, Wa C, DeBolt E, Koke M & Hage DS (2013) Review: Glycation of human serum albumin. Clin Chim Acta 425: 64–76

Bi S, Yan L, Sun Y & Zhang H (2011) Investigation of ketoprofen binding to human serum albumin by spectral methods. Spectrochim Acta Part A Mol Biomol Spectrosc 78: 410– 414

Bohnert T & Gan L-S (2013) Plasma protein binding: from discovery to development. J Pharm Sci 102: 2953–94

Bohnert T, Gan L-S, Howard ML, Hill JJ, Galluppi GR & McLean MA (2010) Plasma protein binding in drug discovery and development. Comb Chem High Throughput Screen 13: 170–87

Borek D, Cymborowski M, Machius M, Minor W & Otwinowski Z (2010) Diffraction data analysis in the presence of radiation damage. Acta Crystallogr D Biol Crystallogr 66: 426–36

Castagna R, Donini S, Colnago P, Serafini A, Parisini E & Bertarelli C (2019) Biohybrid Electrospun Membrane for the Filtration of Ketoprofen Drug from Water. ACS omega 4: 13270–13278

Colclough N, Ruston L, Wood JM & MacFaul PA (2014) Species differences in drug plasma protein binding. Med Chem Commun 5: 963–967

Cooper SA, Reynolds DC, Reynolds B & Hersh E V. (1998) Analgesic Efficacy and Safety of (R)-Ketoprofen in Postoperative Dental Pain. J Clin Pharmacol 38: 11S–18S

Curry S, Mandelkow H, Brick P & Franks N (1998) Crystal structure of human serum albumin complexed with fatty acid reveals an asymmetric distribution of binding sites. Nat Struct Biol 5: 827–35

Czub M, Handing K, Venkataramany BS, Cooper DR, Shabalin IG & Minor W (2020) Albumin-based transport of non-steroidal anti-inflammatory drugs in mammalian blood plasma. J Med Chem

Doweiko JP & Nompleggi DJ (1991) Reviews: Role of Albumin in Human Physiology and Pathophysiology. J Parenter Enter Nutr 15: 207–211

Dubois N, Lapicque F, Abiteboul M & Netter P (1993) Stereoselective protein binding of ketoprofen: Effect of albumin concentration and of the biological system. Chirality 5: 126–134

Emsley P, Lohkamp B, Scott WG & Cowtan K (2010) Features and development of Coot. Acta Crystallogr D Biol Crystallogr 66: 486–501

Ghezzi P, Melillo G, Meazza C, Sacco S, Pellegrini L, Asti C, Porzio S, Marullo A, Sabbatini V, Caselli G, et al (1998) Differential contribution of R and S isomers in ketoprofen anti-inflammatory activity: role of cytokine modulation. J Pharmacol Exp Ther 287: 969–74

Grabowski M, Langner KM, Cymborowski M, Porebski PJ, Sroka P, Zheng H, Cooper DR, Zimmerman MD, Elsliger MA, Burley SK, et al (2016) A public database of macromolecular diffraction experiments. Acta Crystallogr Sect D, Struct Biol 72: 1181– 1193

Hazewinkel HAW, van den Brom WE, Theijse LFH, Pollmeier M & Hanson PD (2003) Reduced dosage of ketoprofen for the short-term and long-term treatment of joint pain in dogs. Vet Rec 152: 11–14

Holm L (2019) Benchmarking fold detection by DaliLite v.5. Bioinformatics 35: 5326–5327

Kantor TG (1986) Ketoprofen: A Review of Its Pharmacologic and Clinical Properties. Pharmacother J Hum Pharmacol Drug Ther 6: 93–102

Ketoprofen entry in Drugbank, https://www.drugbank.ca/drugs/DB01009 [accessed 05.25.2020].

Ketoprofen entry on Drugs.com, https://www.drugs.com/drug-interactions/ketoprofenindex.html [accessed 02.21.2021]

Kosa T, Maruyama T & Otagiri M (1997) Species differences of serum albumins: I. Drug binding sites. Pharm Res 14: 1607–12

Kowiel M, Jaskolski M & Dauter Z (2014) ACHESYM: an algorithm and server for standardized placement of macromolecular models in the unit cell. Acta Crystallogr D Biol Crystallogr 70: 3290–3298

Krissinel E & Henrick K (2007) Inference of macromolecular assemblies from crystalline state. J Mol Biol 372: 774–797

Lees P, Taylor P., Landoni F., Arifah A. & Waters C (2003) Ketoprofen in the Cat: Pharmacodynamics and Chiral Pharmacokinetics. Vet J 165: 21–35

Loet X Le (1989) Safety of Ketoprofen in The Elderly: A Prostective Study on 20,000 Patients. Scand J Rheumatol 18: 21–27

Lombardo F, Berellini G & Obach RS (2018) Trend Analysis of a Database of Intravenous Pharmacokinetic Parameters in Humans for 1352 Drug Compounds. Drug Metab Dispos 46: 1466–1477

Lorier M, Magallanes L, Ibarra M, Guevara N, Vázquez M & Fagiolino P (2016) Stereoselective Pharmacokinetics of Ketoprofen After Oral Administration of Modified-Release Formulations in Caucasian Healthy Subjects. Eur J Drug Metab Pharmacokinet 41: 787–793

Maciążek-Jurczyk M (2014) Phenylbutazone and ketoprofen binding to serum albumin. Fluorescence study. Pharmacol Reports 66: 727–731

Majorek KA, Zimmerman MD, Grabowski M, Shabalin IG, Zheng H & Minor W (2020) Assessment of Crystallographic Structure Quality and Protein–Ligand Complex Structure Validation. In Structural Biology in Drug Discovery pp 253–275. Wiley

Minor W, Cymborowski M, Otwinowski Z & Chruszcz M (2006) HKL-3000: the integration of data reduction and structure solution-from diffraction images to an initial model in minutes. Acta Crystallogr D Biol Crystallogr 62: 859–866

Misra PP & Kishore N (2013) Differential Modulation in Binding of Ketoprofen to Bovine Serum Albumin in the Presence and Absence of Surfactants: Spectroscopic and Calorimetric Insights. Chem Biol Drug Des 82: 81–98

Murshudov GN, Skubák P, Lebedev AA, Pannu NS, Steiner RA, Nicholls RA, Winn MD, Long F & Vagin AA (2011) REFMAC5 for the refinement of macromolecular crystal structures. Acta Crystallogr D Biol Crystallogr 67: 355–367

Otwinowski Z & Minor W (1997) Processing of X-ray diffraction data collected in oscillation mode. Methods Enzymol 276: 307–236

Owens JG, Kamerling SG, Stanton SR & Keowen ML (1995) Effects of ketoprofen and phenylbutazone on chronic hoof pain and lameness in the horse. Equine Vet J 27: 296– 300

Painter J & Merritt EA (2006) Optimal description of a protein structure in terms of multiple groups undergoing TLS motion. Acta Crystallogr Sect D Biol Crystallogr 62: 439–450

Papaneophytou CP, Grigoroudis AI, McInnes C & Kontopidis G (2014) Quantification of the effects of ionic strength, viscosity, and hydrophobicity on protein-ligand binding affinity. ACS Med Chem Lett 5: 931–6

Peters TJT (1995) All About Albumin: Biochemistry, Genetics, and Medical Applications 1st ed. San Diego, CA, USA, CA: Academic Press

Petitpas I, Petersen CE, Ha C, Bhattacharya AA, Zunszain PA, Ghuman J, Bhagavan N V & Curry S (2003) Structural basis of albumin-thyroxine interactions and familial dysalbuminemic hyperthyroxinemia. Proc Natl Acad Sci U S A 100: 6440–6445

Shabalin IG, Porebski PJ & Minor W (2018) Refining the macromolecular model -achieving the best agreement with the data from X-ray diffraction experiment. Crystallogr Rev 24: 236–262

Shen Q, Wang L, Zhou H, Jiang H, Yu L & Zeng S (2013) Stereoselective binding of chiral drugs to plasma proteins. Acta Pharmacol Sin 34: 998–1006

Sudlow G, Birkett DJ & Wade DN (1975) The characterization of two specific drug binding sites on human serum albumin. Mol Pharmacol 11: 824–832

Sudlow G, Birkett DJ & Wade DN (1976) Further characterization of specific drug binding sites on human serum albumin. Mol Pharmacol 12: 1052–1061

Trainor GL (2007) The importance of plasma protein binding in drug discovery. Expert Opin Drug Discov 2: 51–64

Vagin A & Teplyakov A (2010) Molecular replacement with MOLREP. Acta Crystallogr D Biol Crystallogr 66: 22–25

Wang Z, Ho JX, Ruble JR, Rose J, Rüker F, Ellenburg M, Murphy R, Click J, Soistman E, Wilkerson L, et al (2013) Structural studies of several clinically important oncology drugs in complex with human serum albumin. Biochim Biophys Acta 1830: 5356–5374

Williams CJ, Headd JJ, Moriarty NW, Prisant MG, Videau LL, Deis LN, Verma V, Keedy DA, Hintze BJ, Chen VB, et al (2018) MolProbity: More and better reference data for improved all-atom structure validation. Protein Sci 27: 293–315

Winn MD, Ballard CC, Cowtan KD, Dodson EJ, Emsley P, Evans PR, Keegan RM, Krissinel EB, Leslie AGW, McCoy A, et al (2011) Overview of the CCP4 suite and current developments. Acta Crystallogr D Biol Crystallogr 67: 235–242

Zhivkova ZD & Russeva VN (1998) Stereoselective binding of ketoprofen enantiomers to human serum albumin studied by high-performance liquid affinity chromatography. J Chromatogr B Biomed Sci Appl 714: 277–283

Zielinski K, Sekula B, Bujacz A & Szymczak I (2020) Structural investigations of stereoselective profen binding by equine and leporine serum albumins. Chirality 32: 334–344

