## Supplementary Materials for "Transport of Ketoprofen in Mammalian Blood Plasma"

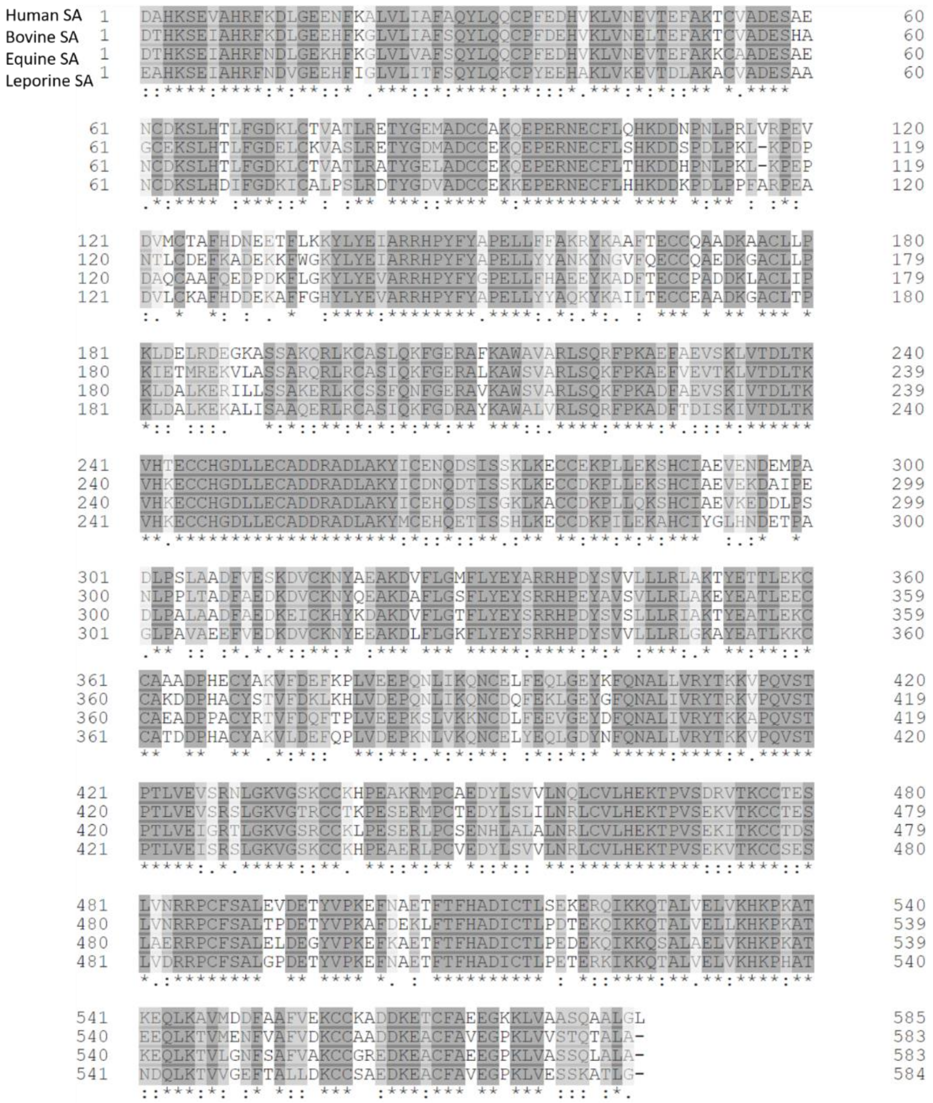


**Figure S1.** Amino acid sequence alignment of human, bovine, equine, and leporine serum albumin. Identical amino acids are labeled in dark gray and with an asterisk. Similar amino acids are labeled with light gray and dots.

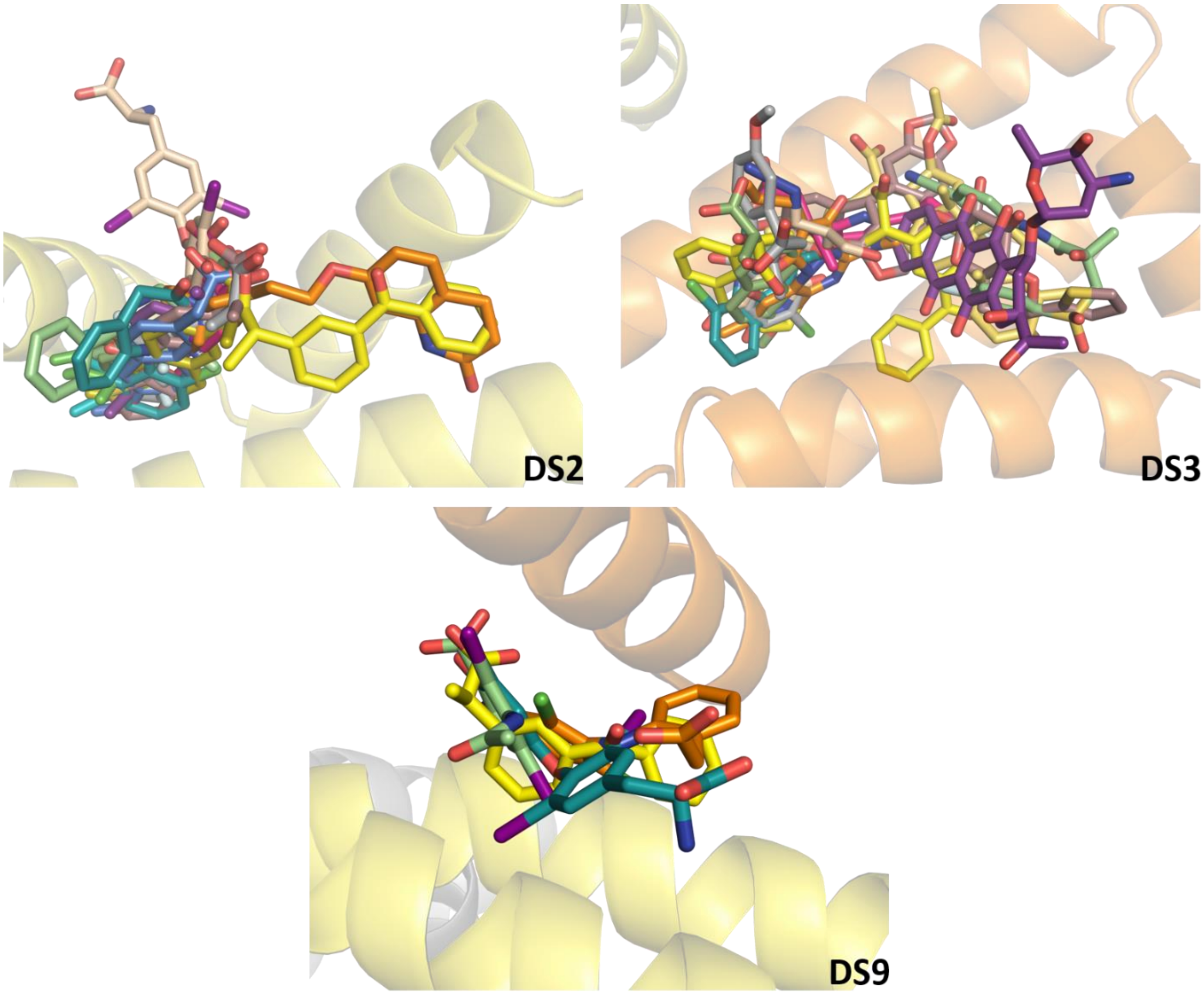


**Figure S2.** Superposition of the HSA-ketoprofen structure (PDB ID: 7JWN) and SA complexes with FDA-approved drugs known to bind to drug sites 2, 3 and 9. Ketoprofen molecules are shown with carbom atoms in yellow on all panels. DS2: aripiprazole (PDB ID: 6A7P), diazepam (PDB ID: 2BXF), diclofenac (PDB ID: 4ZBQ), diflunisal (PDB ID: 2BXE), halothane (PDB ID: 1E7B), ibuprofen (PDB ID: 2BXG), ketoprofen (PDB ID: 6OCK), nabumetone (PDB ID: 6U5A), naproxen (PDB ID: 4ZBR), phenylbutyric acid (PDB ID: 5YOQ), propofol (PDB ID: 1E7A), suprofen (PDB ID: 6OCJ), thyroxine (PDB ID: 1HK1). DS3: azapropazone (PDB ID: 2BXI), bicalutamide (PDB ID: 4LA0), diclofenac (PDB ID: 4Z69), etodolac (PDB ID: 5V0V), fusidic acid (PDB ID: 2VUF), idarubicin (PDB ID: 4LB2), indomethacin (PDB ID: 2BXM), naproxen (PDB ID: 2VDB), salicylic acid (PDB ID: 3B9M), teniposide (PDB ID: 4L9Q), zidovudine (PDB ID: 3B9L). DS9: diclofenac (PDB ID: 6HN0), iodipamine (PDB ID: 2BXN), thyroxine (PDB ID: 1HK4).

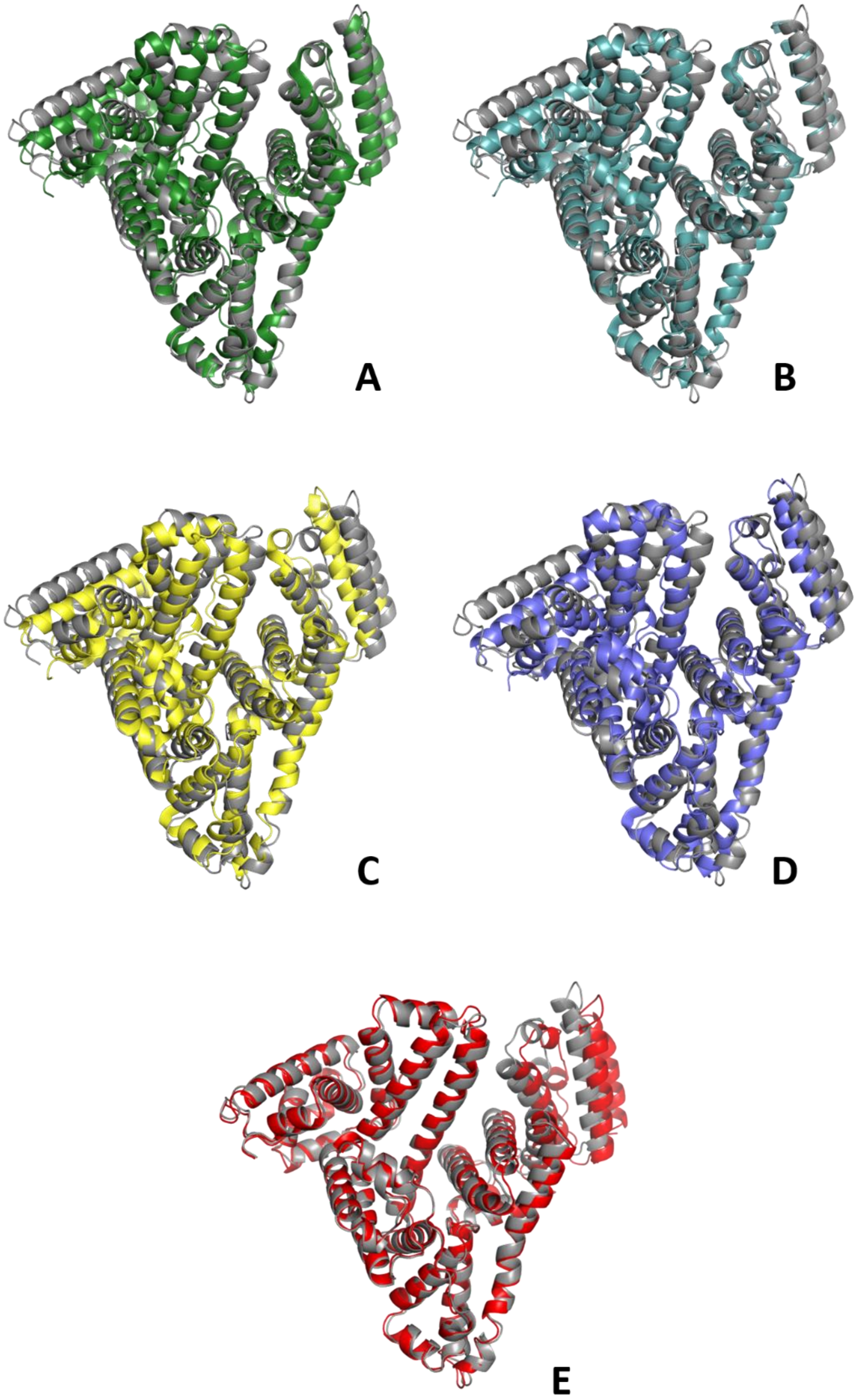


**Figure S3.** Superposition of structure of the HSA-ketoprofen complex (cartoon shown in gray) with the following complexes: A) ESA-ketoprofen (PDB ID: 6U4R); B) BSA-ketoprofen (PDB ID: 6QS9); C) LSA-ketoprofen (PDB ID: 6OCK); D) HSA-ligand free (PDB ID: 4K2C); E) HSA-myristic acid (PDB ID: 1BJ5).

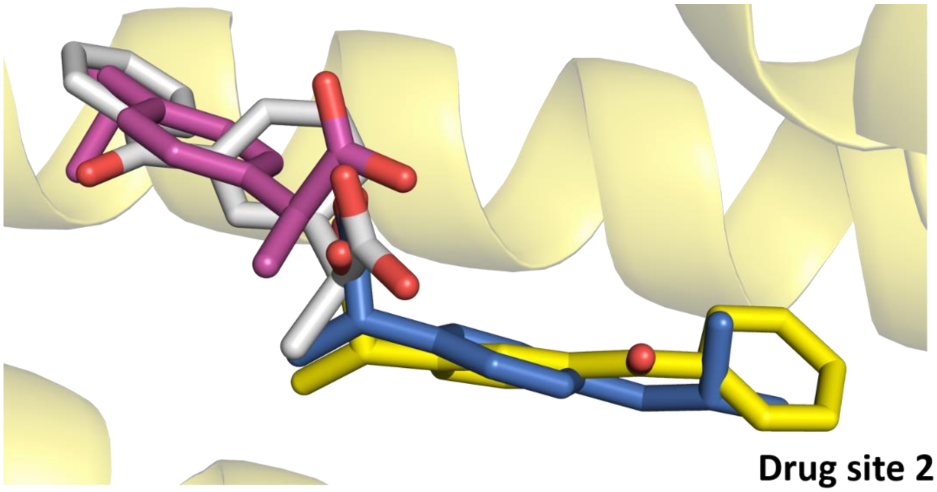


**Figure S4.** Superposition of the crystal structures of ketoprofen complexes with HSA (PDB ID: 7JWN), carbon atoms of the ligand are shown in yellow) and LSA (PDB ID: 6OCK, a ligand in gray) and ibuprofen complexes with HSA (PDB ID: 2BXG, a ligand in pink) and ESA (PDB ID: 6U4X, a ligand in blue).

**Table S1.** RMSD values [Å] between the aligned Cα atoms of SA-ketoprofen complexes, ligand-free SAs, and HSA complex with myristic acid.

| - | HSA-ket (7JWN) | LSA- ket (6OCK) | BSA-ket (6QS9) | ESA-ket (6U4R) | BSA (3V03) | ESA (4F5T) | HSA (4K2C) | LSA (4F5V) | HSA-myr (1BJ5) |
|---|---|---|---|---|---|---|---|---|---|
| **HSA-ket (7JWN)** | - | 4.5 | 4.0 | 3.7 | 4.5 | 3.6 | 3.9 | 4.6 | 1.5 |
| **LSA- ket (6OCK)** | 4.5 | - | 1.5 | 2.4 | 1.5 | 2.3 | 1.8 | 0.7 | 5.2 |
| **BSA- ket (6QS9)** | 4.0 | 1.5 | - | 1.7 | 0.5 | 1.7 | 1.6 | 1.7 | 4.8 |
| **ESA- ket (6U4R)** | 3.7 | 2.4 | 1.7 | - | 1.6 | 0.8 | 1.9 | 2.6 | 3.9 |
| **BSA (3V03)** | 4.5 | 1.5 | 0.5 | 1.6 | - | 1.7 | 1.6 | 1.7 | 4.9 |
| **ESA (4F5T)** | 3.6 | 2.3 | 1.7 | 0.8 | 1.7 | - | 1.8 | 2.5 | 3.9 |
| **HSA (4K2C)** | 3.9 | 1.8 | 1.6 | 1.9 | 1.6 | 1.8 | - | 1.9 | 4.2 |
| **LSA (4F5V)** | 4.6 | 0.7 | 1.7 | 2.6 | 1.7 | 2.5 | 1.9 | - | 5.2 |
| **HSA-myr (1BJ5)** | 1.5 | 5.2 | 4.8 | 3.9 | 4.9 | 3.9 | 4.2 | 5.2 | - |
